# A Workflow to Create Personalised Musculoskeletal Models Based on Magnetic Resonance Images

**DOI:** 10.1101/2025.07.23.666289

**Authors:** Ekaterina Stansfield, Willi Koller, Basílio Gonçalves, Hans Kainz

## Abstract

Musculoskeletal simulations typically rely on generic models that may not accurately represent individual anatomy. While personalisation based on medical images can improve model accuracy, current approaches often require time-consuming workflows to create these models. We present a semi-automatic workflow for creating personalised musculoskeletal models based on magnetic resonance imaging (MRI) that does not require bone segmentation.

Our workflow uses 3D Slicer and Python scripts employing Thin-Plate Spline transformation to map 106 homologous landmarks from generic models onto participants’ anatomy. Generic-scaled and MRI-based models were created for eight healthy participants, and simulations were performed using the participants’ 3D motion capture data. MRI-based models were compared with generic-scaled models through principal component analysis, and joint kinematics and joint contact forces were analysed between both modelling approaches.

Clear geometric differences existed between model types, with MRI-based models showing wider pelvises and different femur/tibia proportions. Unlike generic models, MRI-based male and female models displayed systematic differences. Despite anatomical discrepancies, joint kinematics were similar between models of the same individual, except for pelvis tilt. Muscle moment arms were generally aligned with published data from cadaver studies. MRI-based models consistently produced higher joint contact forces with greater inter-individual variation, particularly at knee joints, compared to generic-scaled models.

The proposed workflow simplifies MRI-based model creation while revealing significant sensitivity of joint contact forces to individual morphology, highlighting the importance of personalisation for biomechanical analyses.

**Author Summary:** Questions about healthy or pathological movement patterns in humans—critical for injury prevention, rehabilitation, and sports performance—are often explored with the help of musculoskeletal modelling. This approach uses a priori defined generic models of the human musculoskeletal system to study joint moments, muscle activation patterns and joint contact forces. Typically, a generic model is scaled to match the participant’s dimensions linearly, which does not allow for an accurate representation of their bone and muscle morphology. We developed a semi-automatic workflow to creating magnetic resonance imaging-based personalisation that does not require bone segmentation but closely matches individual geometry with the help of a non-linear fitting function. By comparing magnetic resonance imaging-based and generic-scaled models in eight individuals, we show systematic bias inherent to one of the most popular musculoskeletal models and demonstrate the importance of model personalisation for healthy adults. Our personalisation pipeline is openly available and easy to set up, which will facilitate musculoskeletal modelling studies based on highly personalised models in clinical and research settings, potentially improving treatment planning and biomechanical assessments in the future.

## Introduction

Musculoskeletal simulations, mostly based on generic-scaled models, are increasingly used to address research questions based on healthy or pathological movement data [1,2]. Such simulations typically require motion data obtained with a 3D gait laboratory. Body segment locations in space are captured using reflective markers placed on the skin. A personalised musculoskeletal model is created either by modifying a pre-existing generic model or by creating a model from scratch [3–5]. Finally, musculoskeletal simulations are performed based on the personalised model and motion data, which typically includes marker trajectories and ground reaction forces [6].

Personalised musculoskeletal models can be created to different degrees, either solely based on the position of reflective markers or by additionally integrating medical images. In ordinary workflows, the position of skin marker pairs is used to define scale factors for personalising a pre-existing generic musculoskeletal model. Based on these scale factors, the model is linearly scaled to match participants’ segment dimensions as closely as possible[7]. However, generic-scaled models do not account for individual anatomical differences, such as variations in bone shape and muscle lines of action [8,9]. To improve accuracy and applicability in medical settings, these models may require further, non-linear personalisation to better align digital representation with actual human subjects.

Integrating information from medical images into model personalisation improves the accuracy of musculoskeletal simulations [10]. The degree of personalisation spans from accurate capture of linear bone measurements to detailed reconstruction of muscle line of action, muscle fibre properties, and joint surface features [5,11,12]. The path to personalised medical imaging-informed models requires several consecutive steps. First, joint centre locations and body segment orientation in parent-child coordinate systems are determined, typically following International Society of Biomechanics (ISB) recommendations [13,14]. Second, correct locations and representations of muscle-tendon paths are identified in response to individual bone morphology. Some approaches to personalise muscle-tendon unit pathways have used linear fitting of muscle via points onto bone surfaces [15], employed non-linear mapping onto bone surfaces [16], utilised segmented muscle surfaces from 3D medical images [12], or applied population-based estimates followed by optimisation algorithms [17,18]. Finally, models’ muscle fibre parameters can be personalised using empirical data from medical images or cadaveric research [19]. Once built, models need validation, for example, by comparing muscle moment arms with published data or estimated muscle activation patterns with recorded electromyography data [20].

The accessibility of open-source segmentation and annotation software, such as 3DSlicer, and open-source musculoskeletal modelling software, such as OpenSim, has made creating personalised musculoskeletal models much more accessible. In this study, we propose a semi-automatic method that utilises Thin-Plate Spline (TPS) function and magnetic resonance imaging-based workflow. A similar methodology for bone reconstruction from sparse data, now known as statistical shape analysis, was previously explored by Zheng et al. [21] and Zheng and Nolte [22]. The present approach, however, is based on identifying biologically and/or geometrically homologous points on template and target objects [23,24]. We compare these MRI-based models with generic-scaled models and investigate the impact of personalisation workflow on inverse kinematics, inverse dynamics and joint contact forces.

## Results

Each of the eight participants (four males and four females, healthy, from 26 to 51 years of age) is represented by two models: a generic-scaled Rajagopal (2016) model that had its linear dimensions scaled to match the participant’s antropometry and an MRI-based model that was obtained with the help of our pipeline (see Methods and Supplementary information).

### Comparing musculoskeletal geometry between generic-scaled and MRI-based models

Bony landmarks and muscle paths of generic-scaled and MRI-based models were analysed using principal components. The first two principal components (PC) described 63.9% of the variance (Fig. 1a). PC1 (51.8% of variance) strongly differentiated between MRI-based and generic-scaled models, ensuring no overlap between groups. PC2 (12.1% of variance) differentiated between MRI-based male and female models. In the generic-scaled models, males and females did not differ.

**Figure 1.**
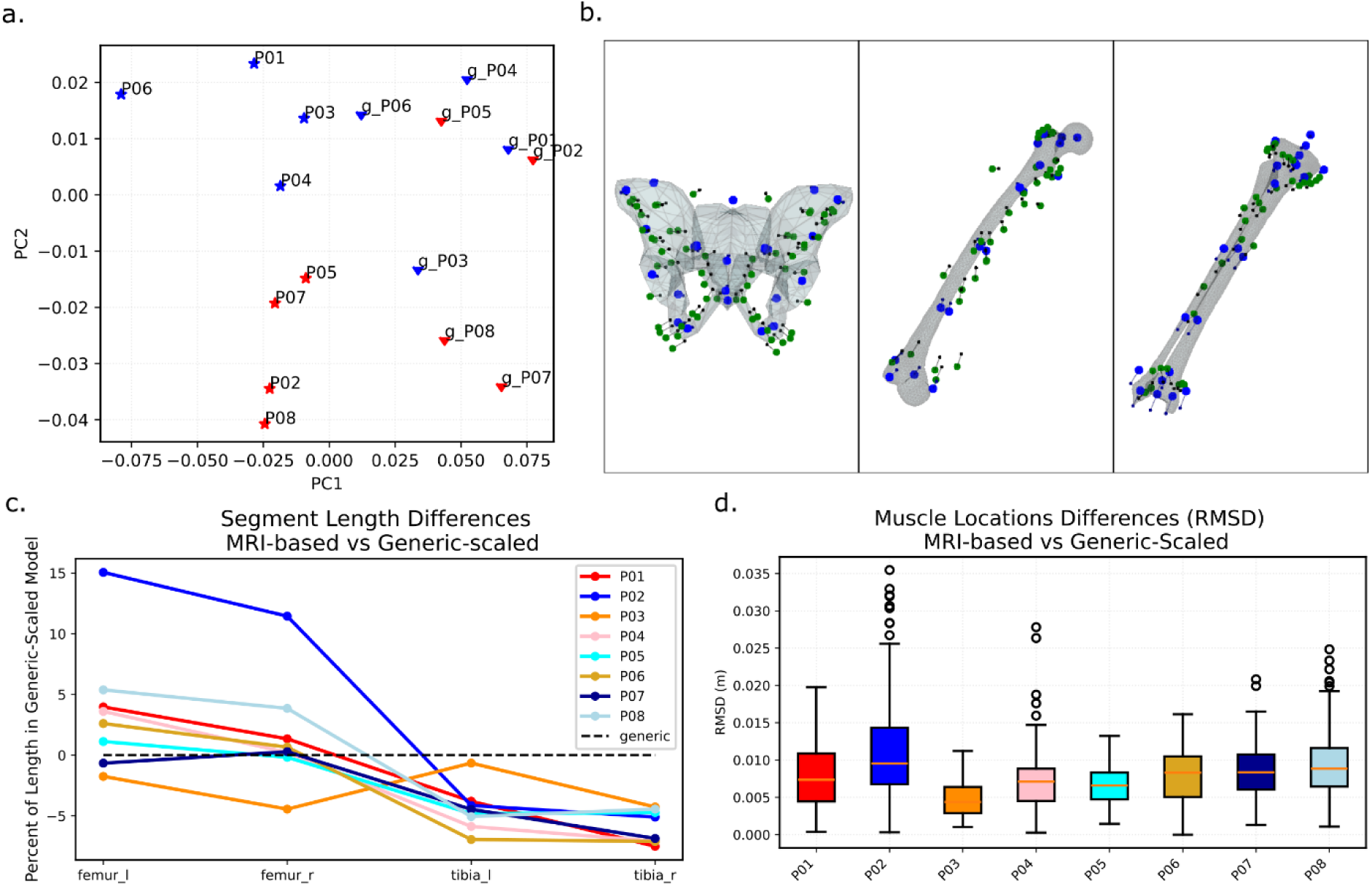
Geometry comparison between MRI-based and generic-scaled models. P01, P02, etc. labels indicate the participant. (a) Principal component analysis of superimposed and standardised data for bone landmarks and muscle path points: the first two principal components; triangles indicate generic-scaled models and stars show MRI-based models. Male models are depicted in red, and female models in blue. (b) Shape differences between generic-scaled and MRI-based models as described by the first principal component. Bone surfaces and small black points indicate bones and muscle paths of an average model across all models. Large blue and green points depict the configuration at PC1 minimum value. Lines between homologous points show direction and magnitude of change between the mean and PC1 minimum. (c) Differences of the segment length in the MRI-based models from the generic-scaled ones, expressed as per cent of the segment length in the generic-based model. (d) Root mean squared distances (RMSD) between generic-scaled and MRI-based models in muscle path point locations.

MRI-based models tend to have wider pelvises (large blue points in Fig. 1b), especially in ischiopubic arches, compared to generic-scaled models, which alters the origins of hamstrings and adductor muscles (large green points in Fig. 1b). In most cases, generic-scaled models underestimate the length of the femur by up to 15% and overestimate the length of the tibia by about 5-7% (Fig. 1c). In articular, femora in the P02 were 12-15% longer in the MRI-based personalised model than in the MRI-based one. Similar observations applied to muscle locations, where P02 had the most extensive distribution of differences in muscle path points, in agreement with segment length differences, and no individual had an average difference of less than 1 cm in muscle via point locations (Fig. 1d).

On further investigation, path point locations for some muscles around the knee in generic-scaled models were notably different compared to their locations in MRIs (Supplementary information, Fig. S2). Muscle origin and via points of *gastrocnemius*, *adductor magnus*, *sartorius,* and *tensor fasciae latae* were located further away from the knee joint centre in generic-scaled models compared to MRI-based models. The sartorius P2 via point was positioned much more anteriorly in the generic-scaled models than its actual location, visible in MRI.

Large errors in muscle insertion points on the patella were observed in the generic-scaled models. Despite scaling the patella to match the participants’ knee widths, the size of the patella in generic-scaled models was significantly larger than the actual size in MRI. As a result, patella muscle via points for *rectus femoris* and *vastii* muscles were at least 1 cm different between generic-scaled and MRI-based models in most individuals (Supplementary information, Fig. S2c).

### Comparing simulation results between generic-scaled and MRI-based models

Joint kinematics was comparable between the generic-scaled and MRI-based models (Fig. 2a). Cosine similarity in joint kinematics between generic-scaled and MRI-based models ranged between 0.8 and 1 (Fig. 2b). Knee and subtalar angles exhibited the least difference between models, and pelvis tilt was affected by the personalisation the most. In inverse dynamic analysis, mean cosine similarity values were consistent for forces and moments across all coordinates.

**Figure 2.**
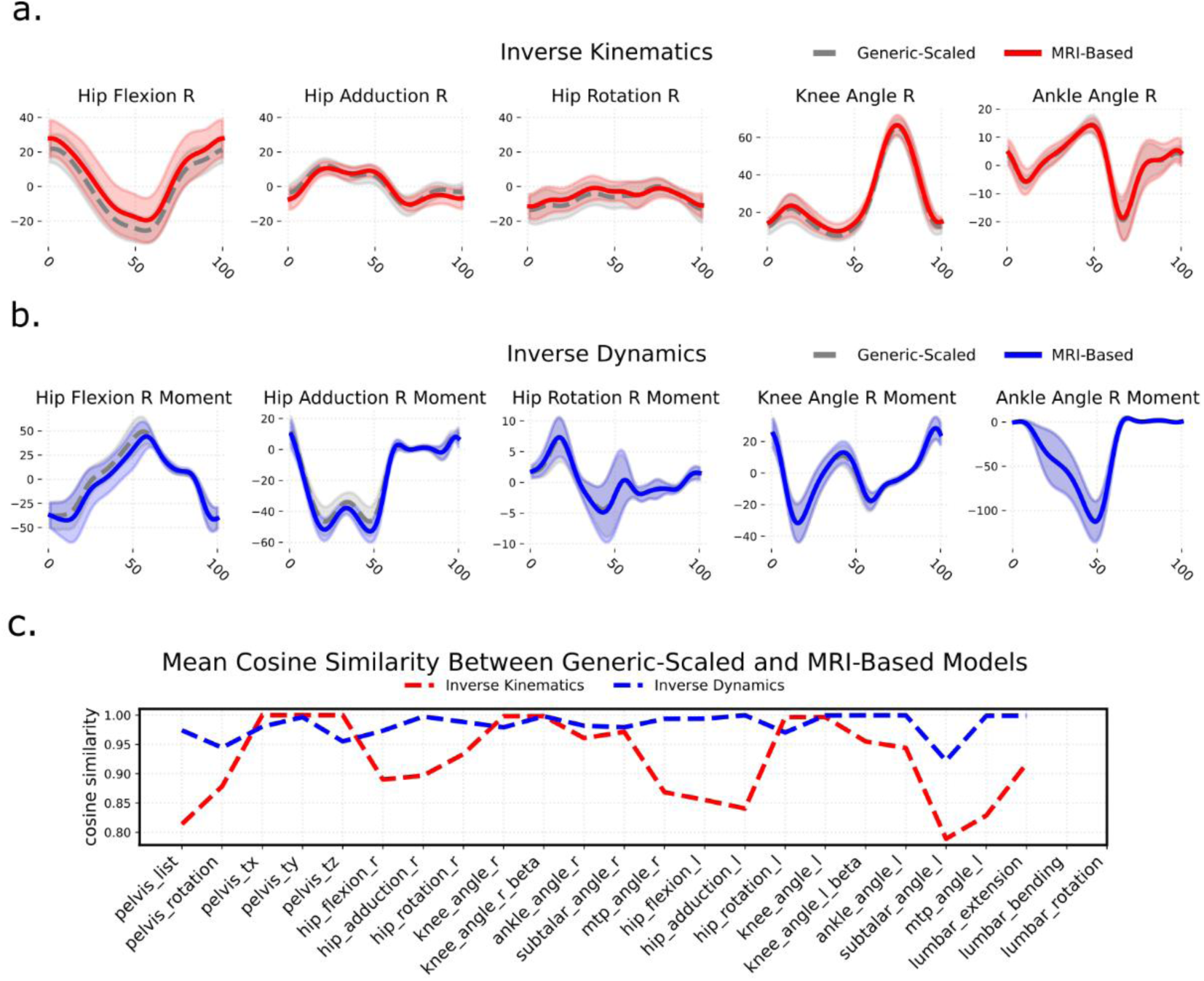
Comparison of kinematics and dynamics between MRI-based and generic-scaled models, averaged across all individuals. (a) Inverse Kinematics, (b) Inverse Dynamics, (c) cosine similarity between mean waveforms in the generic-scaled and MRI-based models.

Geometric differences between models markedly affected muscle moment arms and joint contact forces. The gluteal muscles, *psoas*, and *biceps femoris long head* most frequently showed moment arm differences exceeding 50% of the generic-scaled range across joint angles (Appendix 1, Supplementary information, Tab. S4, Fig. S3). MRI-based models produced higher joint contact forces with greater variation than generic-scaled models (Fig. 3a and b, Supplementary information, Fig. S4). Knees joint contact forces showed the largest increases in median values and the greatest spread among the three joint sets.

**Figure 3.**
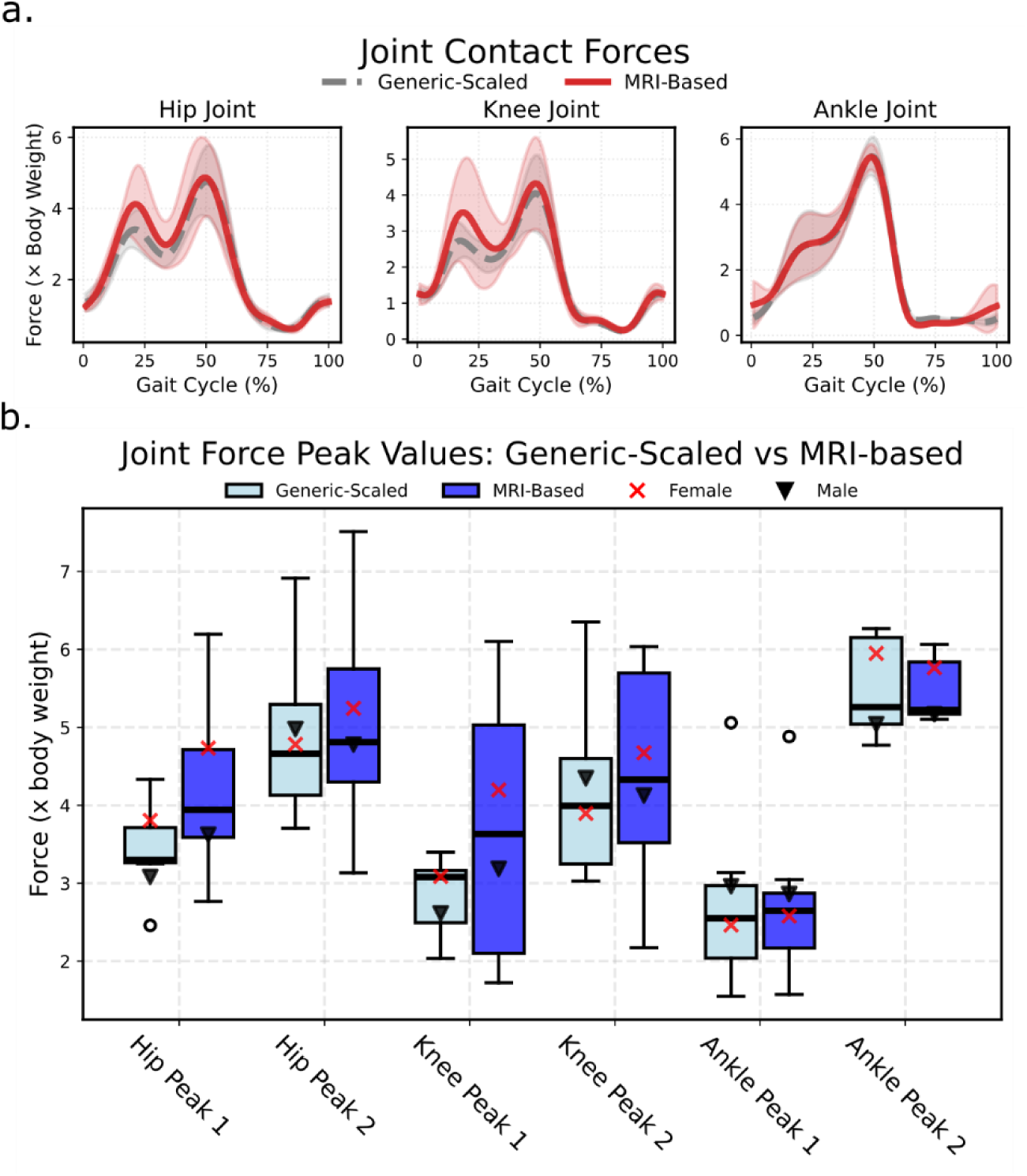
Comparison of joint contact forces calculated with generic-scaled and MRI-based models: (a) Mean waveforms with standard deviation; and (b) boxplot of values exported for first and second peaks of waveforms.

To investigate the relationship between joint contact forces and bone morphology, we computed Pearson’s correlations between peak joint contact forces and the first principal component (PC1), which differentiates between MRI-based and generic-scaled models based on pelvis, femora, patellae, and tibiae shape (Fig. 4a). PC1 exhibited high negative correlations with first peak hip and knee joint contact forces. In other words, generic-scaled models produced lower joint contact forces than their MRI-based counterparts. The same pattern was detected when P07, the individual with the strongest expression of the geometric feature combination as expressed by PC1 (Fig. 1a), was excluded from the correlation test (Fig. 4b).

**Figure 4.**
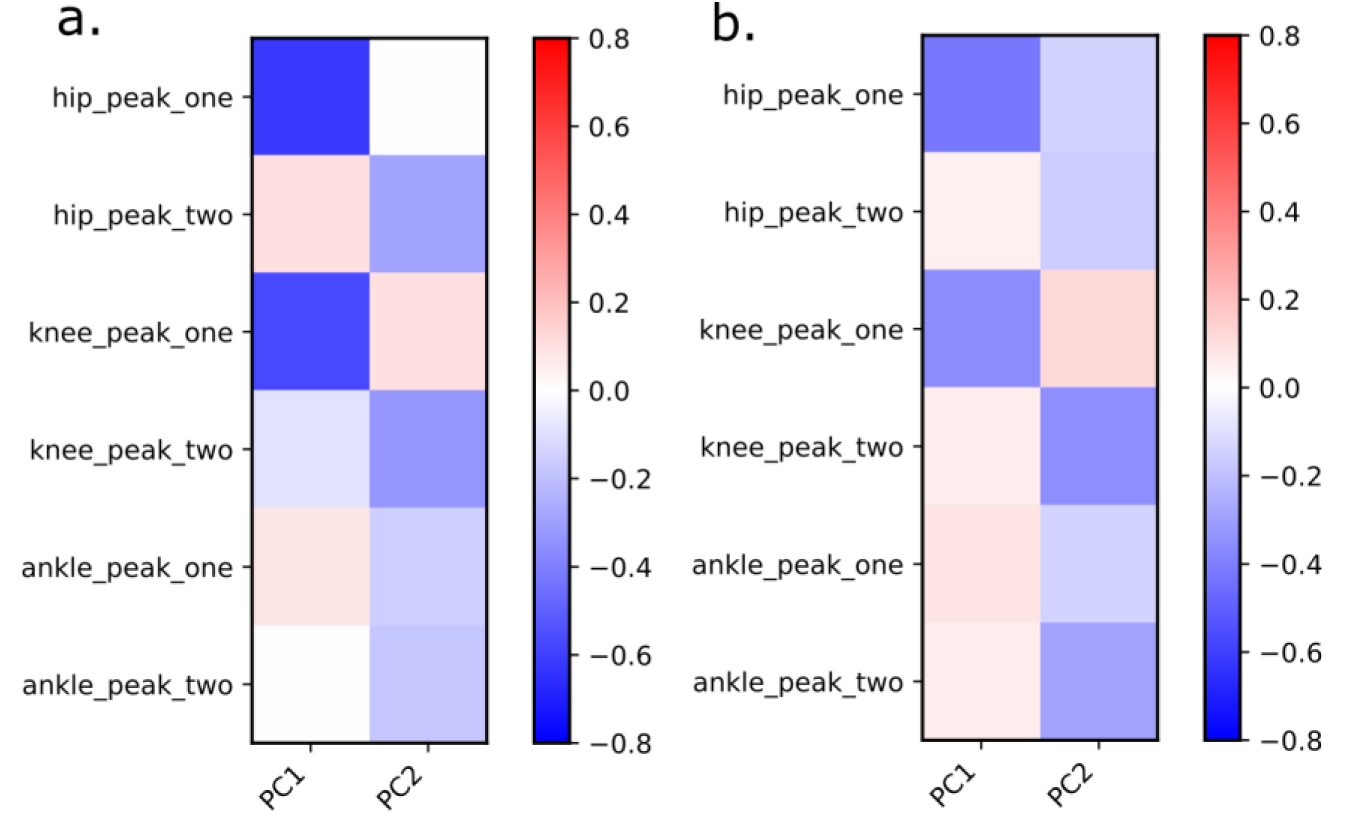
Correlations between the shape of all models (MRI-based and generic-scaled) as described by PC1 peak joint forces: (a) for the complete set of individuals, (b) excluding P07.

### Validation

Inverse kinematics yielded root mean squared (RMS) marker tracking errors of less than 2 cm for all individuals. Across all models, simulations yielded residual forces well below the recommended threshold [20] of 100 N (typically less than 10 N) and residual moments below 35 Nm (typically less than 5 Nm). In most cases, EMG activity corresponds to muscle activations obtained from the MRI-based models (see figures in Appendix 2). However, EMG signal registers earlier than estimated muscle activation, likely due to electromechanical delay between muscle electrical activation (EMG) and mechanical output [30].

Despite discrepancies in leg length and muscle path point locations, muscle moment arms of MRI-based models were, for most muscles, similar to generic-scaled ones (Appendix 1) and, in most cases where literature data were available, corresponded with empirical studies (Supplementary information Fig. S3).

## Discussion

A user-friendly open-source pipeline for a non-linear personalisation of musculoskeletal models was developed using 3D Slicer and custom Python scripts. MRI-based models produced valid kinematics and residual forces output, sensu Hickes et al. [20]. The musculoskeletal geometry of the MRI-based models consistently differed from the generic-scaled models created for the same individual. Nevertheless, muscle moment arms in both generic-scaled and MRI-based models generally fell within two standard deviations of each other and aligned with available empirical data.

Despite anatomical discrepancies, kinematic outputs were broadly comparable between models across most joint coordinates, except for pelvis tilt. This similarity likely results from using identical anatomical coordinate systems and degrees of freedom in both model types [31]. Joint moments showed even greater consistency between models in agreement with previous personalisation studies [32].

Variation in musculoskeletal geometry resulted in differences in muscle moment arms. According to our results, generic-scaled models substantially overestimated patella size, underestimated femur length and overestimated shank length. Consequently, most muscle paths in generic-scaled models deviated from corresponding muscle locations visible in MRIs. The gluteal muscles, *psoas,* and *biceps femoris* were most frequently listed as the muscles with the highest proportional difference between the two models. According to Bosman et al. [33], anatomical variability in muscle via point locations significantly affects calculated muscle forces, with *gluteus medius*, *iliacus*, and *psoas* being the most sensitive. Moreover, perturbations in these muscles’ moment arms may result in extensive concomitant changes in unperturbed muscle forces.

High knee joint contact forces in MRI-based models were associated with higher activations in the *sartorius* and *gastrocnemius* muscles. Additionally, discrepancies in patella size and shape between model types altered via point positions of the *rectus femoris* and *vastii* muscles, reducing their moment arms in MRI-based models. Despite not being maximally activated, these strong muscles contributed substantially to joint contact forces.

Overall, MRI-based models exhibited greater variability in joint contact forces than generic-scaled models, with estimates for some individuals reaching up to 6.5 body weights. Although these values remain within the normal range reported in musculoskeletal modelling studies [10,34,35], they contrast with results from instrumented implants that report forces within 1 to 4 times body weight [36–38].

Correlation analysis revealed that a combination of morphological features in the pelvis, femur, and tibia (PC1) was associated with elevated peak hip and knee contact forces in personalised models. This feature combination was most pronounced in individual P07. Excluding P07 from the analysis maintained the same pattern, although with reduced correlation magnitudes. These results agree with reports that individual variations in femur morphology, such as torsion angle, neck-shaft angle, and femoral neck length, affect joint contact forces [9,39].

Many previous studies have agreed that the physiological consistency of musculoskeletal models improves with the degree of personalisation [15,32,40–44]. Smale et al (2019) and Killen et al. (2024) demonstrated that personalisation of knee geometry results in different kinematics and joint moment patterns, as well as in different knee ligament lengths and joint pressure locations in models for anterior cruciate ligament deficiency patients and knee arthritis. Studies in healthy children and children with cerebral palsy showed a significant impact of personalised musculoskeletal geometry, combined with calibrated parameters of musculotendon units and electromyographic signal tracking, on muscle activation calculations and resulting muscle and joint contact forces [15,32,40–42]. Conconi et al. [32] demonstrated that ankle models with higher complexity, compared to simple hinge models, produced greater adherence to the hypothesis of negligible joint work when quantifying ligament and cartilage deformations from joint motion. Modenese et al. [15] argued that realistic knee contact forces could be estimated when musculotendon parameters were linearly scaled from a reference model and the muscle force-length-velocity relationship was incorporated into the simulation. At the same time, Kainz et al. [40] found that the most realistic joint contact forces were obtained in MRI-based models that tracked electromyography, with geometry personalisation having the greatest impact.

Previous publications in model personalisation, as listed above, frequently focus on children or pathological cases. Here, we present the first demonstration that personalisation also affects outcomes of models created for healthy adults. The personalisation we achieve using a non-linear approach to reconstructing bone morphology and muscle paths is higher than the linear scaling and affine transformation methods adopted in some of the previous studies. For the first time, this approach allowed us to demonstrate systematic differences between male and female musculoskeletal models. These may be responsible for differential muscle activation patterns and joint contact forces between the sexes. A detailed study of a larger population of healthy adults is currently underway.

Our study focuses on musculoskeletal geometry and has several limitations. First, we did not account for individual motor control; EMG-informed approaches may help overcome excessive muscle activations [40]. Second, morphological changes in personalised models may require corresponding muscle–tendon parameter adjustments. Ackland et al. [45] demonstrated that muscle force calculations are highly sensitive to tendon slack length, as altering one muscle’s parameters can influence synergistic muscle forces due to optimisation constraints. Optimisation procedures for adjusting these parameters, as suggested by Modenese et al. [46], may improve outcomes. Finally, we lack a ground truth for joint kinematics and contact forces. However, personalised models produce more accurate results in people with instrumented prostheses [10], making MRI-based models more trustworthy than generic-scaled models.

## Conclusions

Using a non-linear approach, we presented a workflow for MRI-based personalised models that allows for close fitting of skeletal and muscle geometries. We demonstrated that personalisation is essential not only for pathological, paediatric or edge cases, but also for healthy adults. In particular, we clearly showed differences in musculoskeletal geometry between males and females.

Unlike previous work that used non-linear approaches to personalisation, our workflow does not require large population samples to predict muscle paths. It uses open-source tools readily available to both research and medical practitioners. Moreover, our supplementary material includes a step-by-step introduction to creating these models and the required Python code.

Many clinical centres have 3D movement data and medical images but lack the knowledge to create such models. Our proposed workflow may help overcome these limitations and make personalized musculoskeletal modelling more accessible in practice.

## Methods

### Participants

Eight healthy individuals volunteered to participate in this study. Participants’ ages ranged from 26 to 51 (Table 1). They attended a 3D motion capture session at the Department for Biomechanics, Kinesiology and Computer Science in Sport at the University of Vienna (Vienna, Austria), followed by a magnetic resonance imaging (MRI) session at the Orthopaedic Hospital Speising (Vienna, Austria).

**Table 1.** Participants’ statistics, mean (SD)

|  | Number | Age [years] | Mass [kg] | Height [m] |
| --- | --- | --- | --- | --- |
| **Males** | 4 | 26.75 (5.07) | 76.43 (10.32) | 1.84 (0.06) |
| **Females** | 4 | 36.25 (9.15) | 59.53 (5.49) | 1.64 (0.07) |

### Ethics Statement

Ethics approval was obtained from the Ethics Committee of the University of Vienna (reference number 00716). Written informed consent was obtained from all participants prior to data collection.

### Motion Capture

Gait data were collected using a three-dimensional motion capture system (12-camera Vicon Motion Systems, Oxford, UK), five force plates (Kistler Instrumente AG, Winterthur, Switzerland), and a 16-channel electromyography (EMG) system (Cometa SRL, Bareggio, Italy). *Gluteus maximus*, *gluteus medius*, *tensor fasciae latae*, *biceps femoris*, *rectus femoris*, *vastus lateralis*, *vastus medialis* and *gastrocnemius medialis* activation signals were collected on the left and right lower limb with the help of EMG sensors. An adapted Cleveland marker placement protocol (Sutherland, 2002) was used for tracking motion (Supplementary information, Tab. S1).

### MRI images

Each participant underwent MRI scans of the pelvis and lower extremities (Magnetom Vida, Gradient Recalled scanning sequence, segmented k-space variant, Cardiac Gating, Respiratory Gating, Phase Encode Reordering, repetition time 5.42 msec, echo time 2.46 ms, magnetic field strength 3T, imaging frequency 123.25 Hz). Voxel size was 0.9 x 0.9 x 1 mm.

### Model personalisation

We used the Rajagopal et al. [25] model, a commonly used OpenSim model, as a base for creating personalised models. The arms were removed from the base model, and the torso mass was adjusted to compensate for the loss of the arms. A generic-scaled and MRI-based models were created for each participant. MRI-based models featured joint centres in MRI according to ISB definitions and muscle paths closely matching actual anatomy. Bone surfaces were approximated for display purposes only.

#### Generic-scaled models

The generic Rajagopal model was scaled to match participant dimensions determined by reflective markers during static motion capture (Supplementary information, Tab. S2). Scaling was performed using the OpenSim scaling tool, which linearly updates x, y, and z segment dimensions and scales muscle fibres’ optimal length to individual segment dimensions. We also adjusted maximum isometric muscle force using a regression equation from Handsfield et al. [26] for calculating lower limb muscle volumes. The maximum muscle force scaling factor was determined by equation (1).

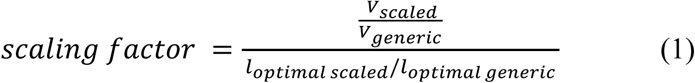

Where 𝑉_𝑠𝑐𝑎𝑙𝑒𝑑_, 𝑉_𝑔𝑒𝑛𝑒𝑟𝑖𝑐_, are volumes of the generic and scaled models, 𝑙_𝑜𝑝𝑡𝑖𝑚𝑎𝑙 𝑠𝑐𝑎𝑙𝑒𝑑_ and 𝑙_𝑜𝑝𝑡𝑖𝑚𝑎𝑙 𝑔𝑒𝑛𝑒𝑟𝑖𝑐_ are optimal fibre lengths of a specific muscle in the scaled individual and generic models. In their turn, the volumes were defined by equation (2) [26].

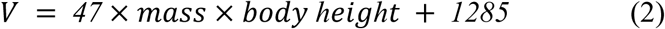

This equation agrees with the convention that maximum isometric muscle force is proportional to physiological cross-sectional area (PCSA). The scaled-generic model, together with muscle parameters and inertia values, became the template for creating the MRI-based models for each participant.

#### MRI-based models

A workflow to create the MRI-based model was developed using 3DSlicer software and OpenSim API in Python (Supplementary information, Fig. S1). Creating the MRI-based model included updating the generic-scaled model with muscle paths and bone shapes obtained from MRI images. To obtain these data, we identified homologous landmarks on the generic model and participants’ MRIs. These landmarks were chosen to represent each segment’s geometry while providing sufficient guidance for later muscle path fitting. 106 landmarks were placed on the pelvis, femur, patella, tibia and fibula of the template model (Fig. 5, Supplementary information Tab. S3). Landmarks represented two groups. The primary group included 21 landmarks easily identifiable on MRI and essential for further steps, placed manually on MRI using 3DSlicer (Fig. 5). Remaining landmarks were projected into MRI space using a Thin-Plate Spline (TPS) non-linear registration algorithm available in 3DSlicer. An operator verified the correct positioning of registered landmarks to ensure alignment with predefined orientation planes within the MRI volume. Once proper positioning was established, bone landmarks were used for Thin-Plate Spline fitting of muscle point paths and translations of muscle wrapping surfaces.

**Figure 5.**
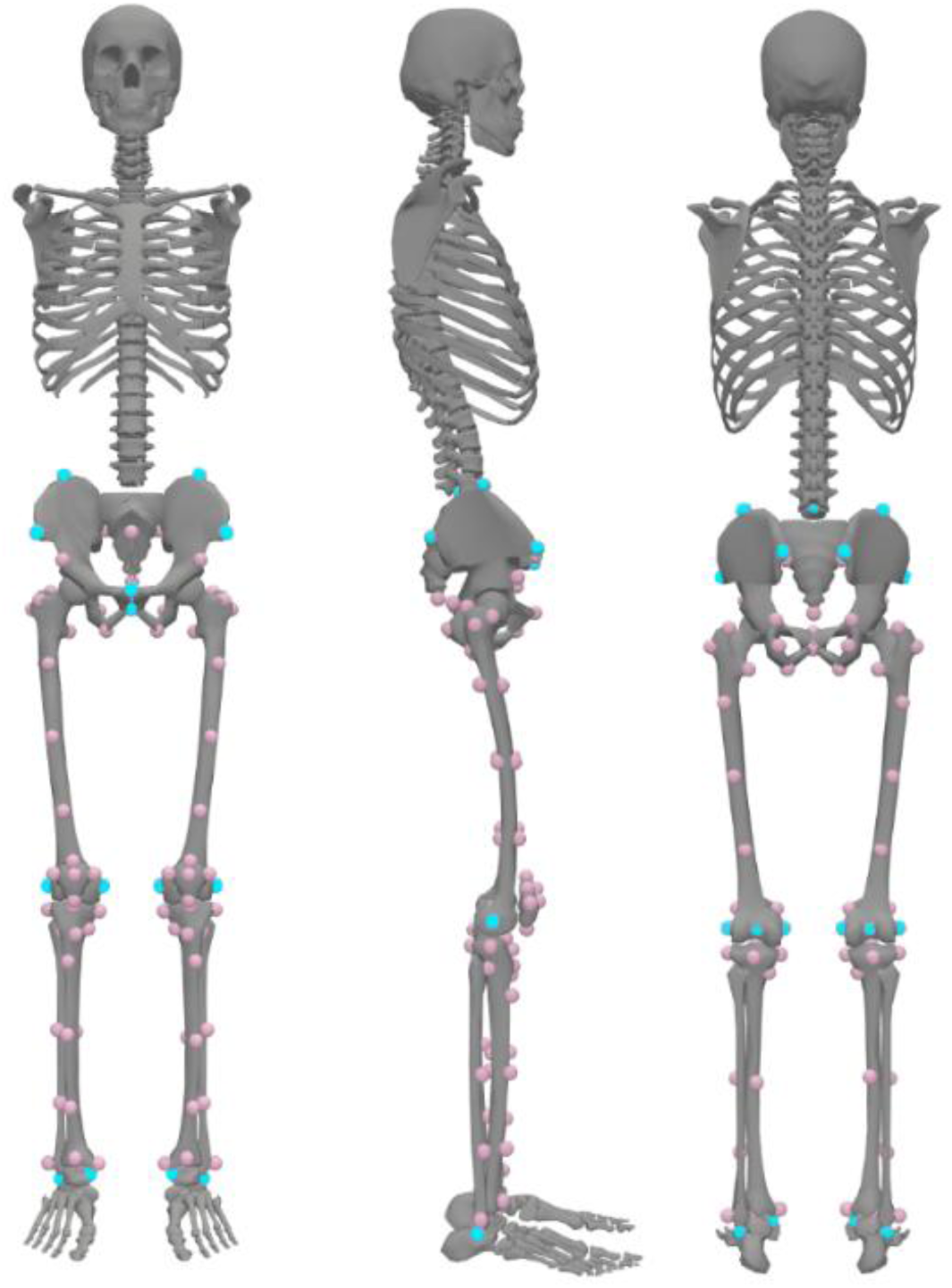
Locations of homologous landmarks on the generic model. The primary set of 21 landmarks is shown in blue. Landmarks in femur head centres are not visible. The secondary set comprising 89 landmarks is shown in pink.

Fitting was achieved using a custom Python script, though the same procedure can be set up using only 3DSlicer tools. Thin-Plate Spline bending strength was penalised at a 0.02 level, determined experimentally to ensure projected points fell within the expected boundaries of bone and muscle positions. Once projected, muscle path locations were rechecked to fit in the middle of the muscles’ axial cross-sections in MRI space. Only a few manual adjustments were necessary (Fig. 6).

**Figure 6.**
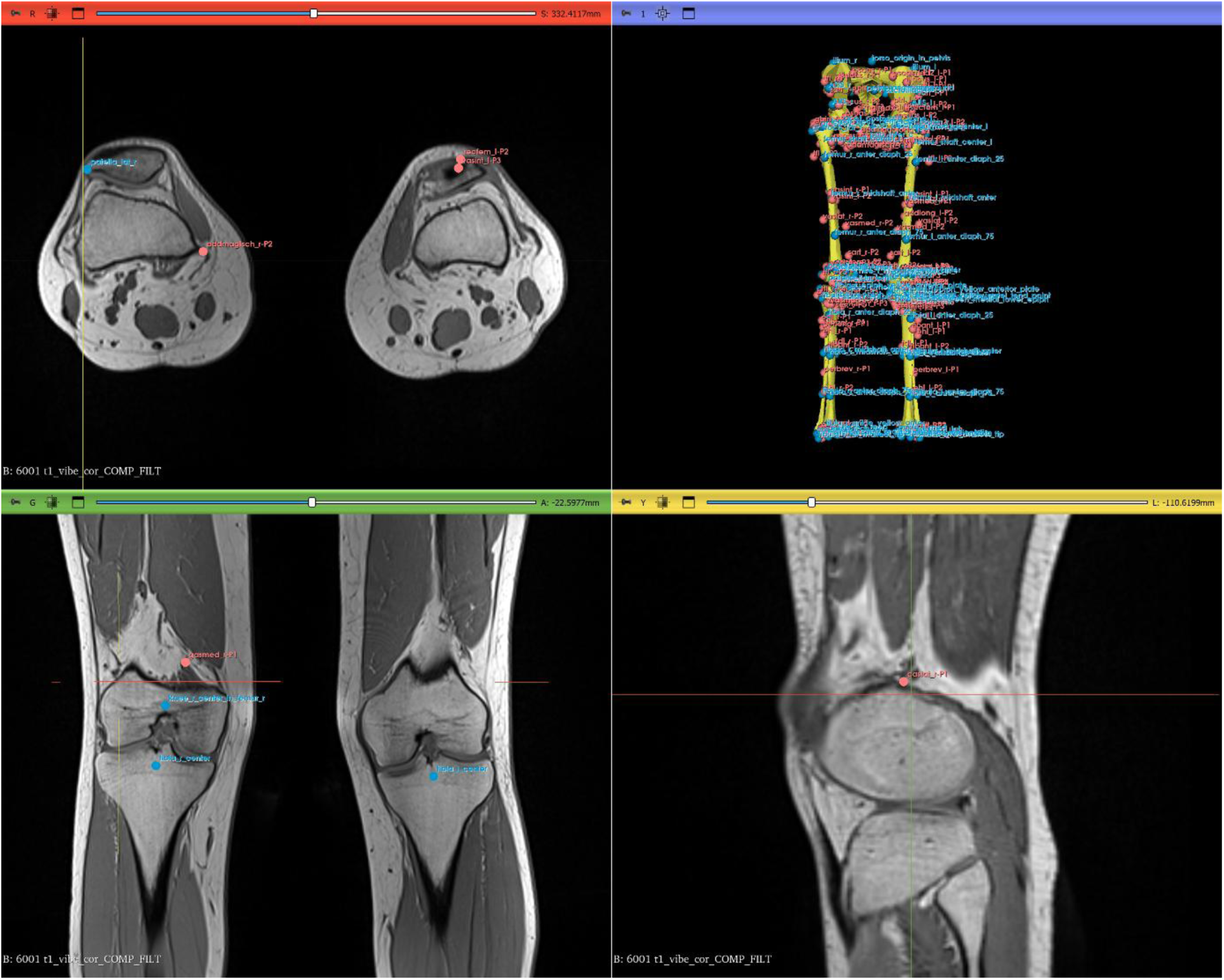
Landmarks in 3D Slicer before data transformation into OpenSim coordinate sets. Blue points show bony landmarks. Pink points show muscle paths. Yellow bone representations were created by warping the template OpenSim geometry onto the participant’s landmark data.

Next, the data were separated into segments, and each segment was rotated to ISB-recommended orientation and scaled to meters to match OpenSim scaling convention. Simultaneously, new bone surfaces were created by warping template model bone surfaces onto participant data. Surfaces were rotated, translated into the ISB-recommended orientation, and later used for model display.

Once muscle paths, joint locations, muscle wrapping surface translations and bone surfaces were obtained and transformed to ISB standard orientation, a copy of the generic-scaled model was updated with this information. Wrapping surface radii, length, and orientation were preserved from the generic-scaled model. During adjustment, a custom Python script monitored potential intersections between muscle path points and wrapping surfaces to ensure accurate alignment. The wrapping surface radius was modified if the distance of the neighbouring via point from the axis of the wrapping surface was smaller than its radius. Skin marker locations underwent TPS transformation to fit the personalised geometry, after which experimental marker locations from static trials were fitted using the OpenSim scaling tool.

Personalisation frequently results in discrepancies between muscle paths and wrapping surface size and position throughout the entire range of motion [27]. To address this, we created an additional optimisation script that adjusted muscle via point positions to prevent muscles from dynamically penetrating wrapping surfaces and avoid significant decreases in personalised muscle moment arms. The following objective function was minimised:

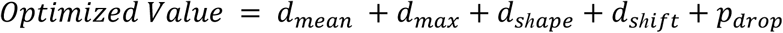

where 𝑑_𝑚𝑒𝑎𝑛_ means the average absolute distance between the generic and personalised muscle moment arm; 𝑑_𝑚𝑎𝑥_ means the maximum absolute distance between the moment arms; 𝑑_𝑠*ℎ*𝑎𝑝𝑒_ is the term that calculates the absolute maximum height between the start and the end of the moment arm waveform; and 𝑝_𝑑𝑟𝑜𝑝_ is the penalty for the sudden drop in the moment arm due to the interaction with the wrapping surface. We utilised an OpenSim API function calculating moment arm as the differential of the ratio between musculotendon length and joint angle [28]. Adjustment boundaries were customised for different muscles.

For example, *gluteus maximus* via points could move within a 2 cm radius, while *rectus femoris* and *vastii* points at the patella were limited to a 0.5 cm radius. The optimisation procedure included grid search within the set radius, followed by gradient optimisation using the Sequential Least Squares Programming (SLSQP) method in the SciPy Python library.

Python scripts and notebooks required to reproduce this pipeline are available publicly.

### Validation and Analyses

#### Comparing generic-scaled and MRI-based models

To compare the musculoskeletal geometry of the generic-scaled and MRI-based models, we first extracted principal components from the clouds of landmarks on the bone and muscle path points, which combined data from all models. The components were extracted from the superimposed and standardised landmark clouds, so variations due to translation, orientation, and size were removed from the data [29]. This allowed us to explore the trends in the shape differences between the generic-scaled and MRI-based models. We also computed and plotted root mean square differences (RMSD) in joint and muscle locations between generic-scaled and MRI-based models.

Inverse kinematics, muscle moment arm analyses, inverse dynamics, static optimisation, and joint reaction analyses were performed using custom Python scripts and the OpenSim 4.5 API. Joint kinematics and dynamics were compared between the MRI-based and generic-scaled models. This was achieved by calculating the cosine similarity metric, a cos between two vectors of values for joint angles (in kinematics) or joint moments (in dynamics) for the right gait cycle in each individual. We further compared predicted joint contact forces in generic-scaled models with MRI-based models to quantify the sensitivity of joint contact forces to model personalisation.

#### Validation

We compared kinematic tracking errors and residual forces with recommended threshold values [20]. Furthermore, model validation was performed by comparing muscle activations estimated by static optimisation with EMG data collected from 16 muscles. Additionally, we compared muscle moment arms from generic-scaled and MRI-based models with data from the literature.

## Acknowledgement

ES acknowledges funding by the Austrian Science Fund, grant P 35714-B. HK and WK acknowledge funding by the European Research Council, GAP 101170218. Co-funded by the European Union. Views and opinions expressed are however those of the authors only and do not necessarily reflect those of the European Union or European Research Council Executive Agency. Neither the European Union nor the granting authority can be held responsible for them.

## Data availability

Scripts for re-creating the pipeline, models and motion capture data are available at https://github.com/katya-stanzy/tps_personalised_model.

## References

[1] Buehler C, Koller W, De Comtes F, Kainz H. Quantifying muscle forces and joint loading during hip exercises performed with and without an elastic resistance band. Front Sports Act Living 2021;3:695383.

[2] Van Rossom S, Kainz H, Wesseling M, Papageorgiou E, De Groote F, Van Campenhout A, et al. Single-event multilevel surgery, but not botulinum toxin injections normalize joint loading in cerebral palsy patients. Clin Biomech (Bristol, Avon) 2020;76:105025.

[3] Kainz H, Hoang HX, Stockton C, Boyd RR, Lloyd DG, Carty CP. Accuracy and Reliability of Marker-Based Approaches to Scale the Pelvis, Thigh, and Shank Segments in Musculoskeletal Models. J Appl Biomech 2017;33:354–60.

[4] Scheys L, Van Campenhout A, Spaepen A, Suetens P, Jonkers I. Personalized MR-based musculoskeletal models compared to rescaled generic models in the presence of increased femoral anteversion: effect on hip moment arm lengths. Gait Posture 2008;28:358–65.

[5] Valente G, Crimi G, Vanella N, Schileo E, Taddei F. nmsBuilder: Freeware to create subject-specific musculoskeletal models for OpenSim. Comput Methods Programs Biomed 2017;152:85–92.

[6] Seth A, Sherman M, Reinbolt JA, Delp SL. OpenSim: a musculoskeletal modeling and simulation framework for in silico investigations and exchange. Procedia IUTAM 2011;2:212–32.

[7] Delp SL, Anderson FC, Arnold AS, Loan P, Habib A, John CT, et al. OpenSim: open-source software to create and analyze dynamic simulations of movement. IEEE Trans Biomed Eng 2007;54:1940–50.

[8] Kainz H, Jonkers I. Imaging-based musculoskeletal models alter muscle and joint contact forces but do not improve the agreement with experimentally measured electromyography signals in children with cerebral palsy. Gait Posture 2023;100:91–5.

[9] Kainz H, Mindler GT, Kranzl A. Influence of femoral anteversion angle and neck-shaft angle on muscle forces and joint loading during walking. PLoS One 2023;18:e0291458.

[10] Modenese L, Barzan M, Carty CP. Dependency of lower limb joint reaction forces on femoral version. Gait Posture 2021;88:318–21.

[11] Scheys L, Desloovere K, Spaepen A, Suetens P, Jonkers I. Calculating gait kinematics using MR-based kinematic models. Gait Posture 2011;33:158–64.

[12] Modenese L, Kohout J. Automated Generation of Three-Dimensional Complex Muscle Geometries for Use in Personalised Musculoskeletal Models. Ann Biomed Eng 2020;48:1793–804.

[13] Wu G, Siegler S, Allard P, Kirtley C, Leardini A, Rosenbaum D, et al. ISB recommendation on definitions of joint coordinate system of various joints for the reporting of human joint motion— part I: ankle, hip, and spine. J Biomech 2002;35:543–8.

[14] Wu G, van der Helm FCT, Veeger HEJD, Makhsous M, Van Roy P, Anglin C, et al. ISB recommendation on definitions of joint coordinate systems of various joints for the reporting of human joint motion--Part II: shoulder, elbow, wrist and hand. J Biomech 2005;38:981–92.

[15] Modenese L, Montefiori E, Wang A, Wesarg S, Viceconti M, Mazzà C. Investigation of the dependence of joint contact forces on musculotendon parameters using a codified workflow for image-based modelling. J Biomech 2018;73:108–18.

[16] Scheys L, Jonkers I, Loeckx D, Maes F, Spaepen A, Suetens P. Image Based Musculoskeletal Modeling Allows Personalized Biomechanical Analysis of Gait. Biomedical Simulation, Springer Berlin Heidelberg; 2006, p. 58–66.

[17] Killen BA, Brito da Luz S, Lloyd DG, Carleton AD, Zhang J, Besier TF, et al. Automated creation and tuning of personalised muscle paths for OpenSim musculoskeletal models of the knee joint. Biomech Model Mechanobiol 2021;20:521–33.

[18] Nolte D, Tsang CK, Zhang KY, Ding Z, Kedgley AE, Bull AMJ. Non-linear scaling of a musculoskeletal model of the lower limb using statistical shape models. J Biomech 2016;49:3576–81.

[19] Bolsterlee B, Veeger HEJD, van der Helm FCT, Gandevia SC, Herbert RD. Comparison of measurements of medial gastrocnemius architectural parameters from ultrasound and diffusion tensor images. J Biomech 2015;48:1133–40.

[20] Hicks JL, Uchida TK, Seth A, Rajagopal A, Delp SL. Is my model good enough? Best practices for verification and validation of musculoskeletal models and simulations of movement. J Biomech Eng 2015;137:020905.

[21] Zheng G, Rajamani KT, Nolte L-P. Use of a dense surface point distribution model in a three-stage anatomical shape reconstruction from sparse information for computer assisted orthopaedic surgery: A preliminary study. Computer Vision – ACCV 2006, Berlin, Heidelberg: Springer Berlin Heidelberg; 2006, p. 52–60.

[22] Zheng G, Nolte L-P. Surface reconstruction of bone from X-ray images and point distribution model incorporating a novel method for 2D-3D correspondence. 2006 IEEE Computer Society Conference on Computer Vision and Pattern Recognition - Volume 2 (CVPR’06), IEEE; 2006. 10.1109/cvpr.2006.300.

[23] Bookstein FL. Thin-Plate splines and the atlas problem for biomedical images. In: Colchester ACF, Hawkes DJ, editors. Information Processing in Medical Imaging, Berlin/Heidelberg: Springer-Verlag; 1991, p. 326–42.

[24] Slice DE. Modern morphometrics. Modern Morphometrics in Physical Anthropology 2005.

[25] Rajagopal A, Dembia CL, DeMers MS, Delp DD, Hicks JL, Delp SL. Full-Body Musculoskeletal Model for Muscle-Driven Simulation of Human Gait. IEEE Trans Biomed Eng 2016;63:2068–79.

[26] Handsfield GG, Meyer CH, Hart JM, Abel MF, Blemker SS. Relationships of 35 lower limb muscles to height and body mass quantified using MRI. Journal of Biomechanics 2014;47:631–8.

[27] Koller W, Horsak B, Kranzl A, Unglaube F, Baca A, Kainz H. Physiological plausible muscle paths: A MATLAB tool for detecting and resolving muscle path discontinuities in musculoskeletal OpenSim models. Gait Posture 2025;117:S21–2.

[28] Sherman MA, Seth A, Delp SL. WHAT IS A MOMENT ARM? CALCULATING MUSCLE EFFECTIVENESS IN BIOMECHANICAL MODELS USING GENERALIZED COORDINATES. Proc ASME Des Eng Tech Conf 2013;2013. 10.1115/DETC2013-13633.

[29] Dryden IL, Mardia KV. Statistical Shape Analysis: With Applications in R. John Wiley & Sons; 2016.

[30] Cavanagh PR, Komi PV. Electromechanical delay in human skeletal muscle under concentric and eccentric contractions. Eur J Appl Physiol Occup Physiol 1979;42:159–63.

[31] Kainz H, Modenese L, Lloyd DG, Maine S, Walsh HPJ, Carty CP. Joint kinematic calculation based on clinical direct kinematic versus inverse kinematic gait models. J Biomech 2016;49:1658–69.

[32] Conconi M, Montefiori E, Sancisi N, Mazzà C. Modeling musculoskeletal dynamics during gait: Evaluating the best personalization strategy through model anatomical consistency. Appl Sci (Basel) 2021;11:8348.

[33] Bosmans L, Valente G, Wesseling M, Van Campen A, De Groote F, De Schutter J, et al. Sensitivity of predicted muscle forces during gait to anatomical variability in musculotendon geometry. J Biomech 2015;48:2116–23.

[34] Giarmatzis G, Jonkers I, Wesseling M, Van Rossom S, Verschueren S. Loading of hip measured by hip contact forces at different speeds of walking and running: Hip loading measured by hcfs at different speeds of walking and running. J Bone Miner Res 2015;30:1431–40.

[35] Kainz H, Killen BA, Wesseling M, Perez-Boerema F. A multi-scale modelling framework combining musculoskeletal rigid-body simulations with adaptive finite element analyses, to evaluate the impact of femoral geometry …. PLoS 2020.

[36] Stansfield BW, Nicol AC, Paul JP, Kelly IG, Graichen F, Bergmann G. Direct comparison of calculated hip joint contact forces with those measured using instrumented implants. An evaluation of a three-dimensional mathematical model of the lower limb. J Biomech 2003;36:929–36.

[37] Bergmann G, Bender A, Dymke J, Duda G, Damm P. Standardized loads acting in hip implants. PLoS One 2016;11:e0155612.

[38] Damm P, Kutzner I, Bergmann G, Rohlmann A, Schmidt H. Comparison of in vivo measured loads in knee, hip and spinal implants during level walking. J Biomech 2017;51:128–32.

[39] Lenaerts G, De Groote F, Demeulenaere B, Mulier M, Van der Perre G, Spaepen A, et al. Subject-specific hip geometry affects predicted hip joint contact forces during gait. J Biomech 2008;41:1243–52.

[40] Kainz H, Wesseling M, Jonkers I. Generic scaled versus subject-specific models for the calculation of musculoskeletal loading in cerebral palsy gait: Effect of personalized musculoskeletal …. Clin Biomech 2021.

[41] Davico G, Lloyd DG, Carty CP, Killen BA, Devaprakash D, Pizzolato C. Multi-level personalization of neuromusculoskeletal models to estimate physiologically plausible knee joint contact forces in children. Biomech Model Mechanobiol 2022;21:1873–86.

[42] Davico G, Pizzolato C, Lloyd DG, Obst SJ, Walsh HPJ, Carty CP. Increasing level of neuromusculoskeletal model personalisation to investigate joint contact forces in cerebral palsy: A twin case study. Clin Biomech (Bristol, Avon) 2020;72:141–9.

[43] Killen BA, Willems M, Jonkers I. An open-source framework for the generation of OpenSim models with personalised knee joint geometries for the estimation of articular contact mechanics. J Biomech 2024;177:112387.

[44] Smale KB, Conconi M, Sancisi N, Krogsgaard M, Alkjaer T, Parenti-Castelli V, et al. Effect of implementing magnetic resonance imaging for patient-specific OpenSim models on lower-body kinematics and knee ligament lengths. J Biomech 2019;83:9–15.

[45] Ackland DC, Lin Y-C, Pandy MG. Sensitivity of model predictions of muscle function to changes in moment arms and muscle-tendon properties: a Monte-Carlo analysis. J Biomech 2012;45:1463–71.

[46] Modenese L, Ceseracciu E, Reggiani M, Lloyd DG. Estimation of musculotendon parameters for scaled and subject specific musculoskeletal models using an optimization technique. J Biomech 2016;49:141–8.

